## Supplemental Data for "Sulfation affects apical extracellular matrix organization during development of the *Drosophila* embryonic salivary gland tube"

### Figure 1-figure supplement 1

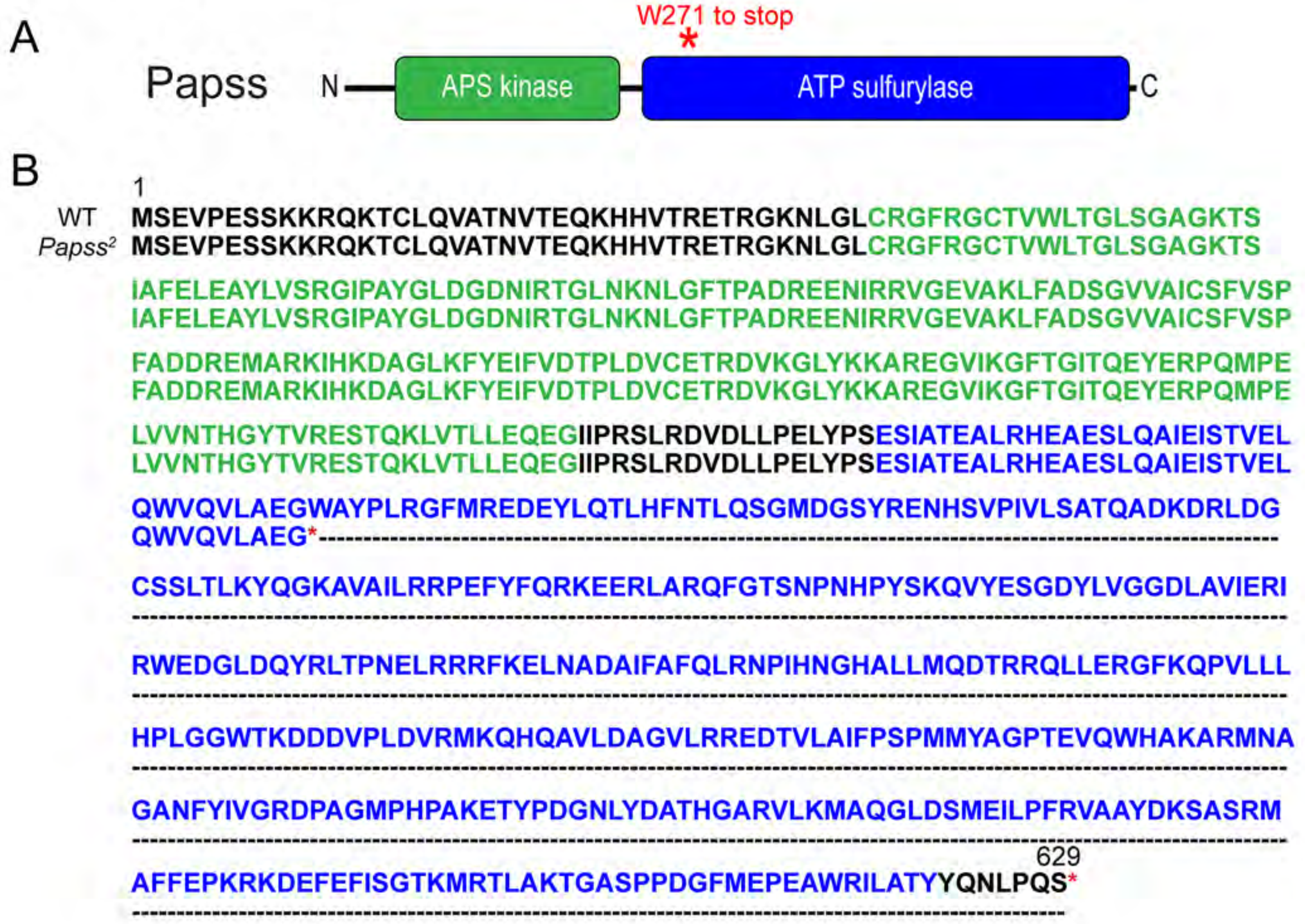


**Figure 1-figure supplement 1. *Papss* mutants encode a stop codon in the ATP sulfurylase domain.** (A) Cartoon of the Papss-PA isoform showing the APS kinase and ATP sulfurylase domains. A red asterisk indicates the site of the W271->stop mutation. (B) Amino acid sequences for WT and *Papss*$^{2}$ sequences for the Papss-PA isoform. Green text, residues of the APS kinase domain. Blue text, residues of the ATP sulfurase domain. Red asterisks, stop codon. Note that *Papss*$^{2}$ also has a secondary mutation (G->A) at G79 of the transcript resulting in a silent mutation.

Figure 1-figure supplement 2

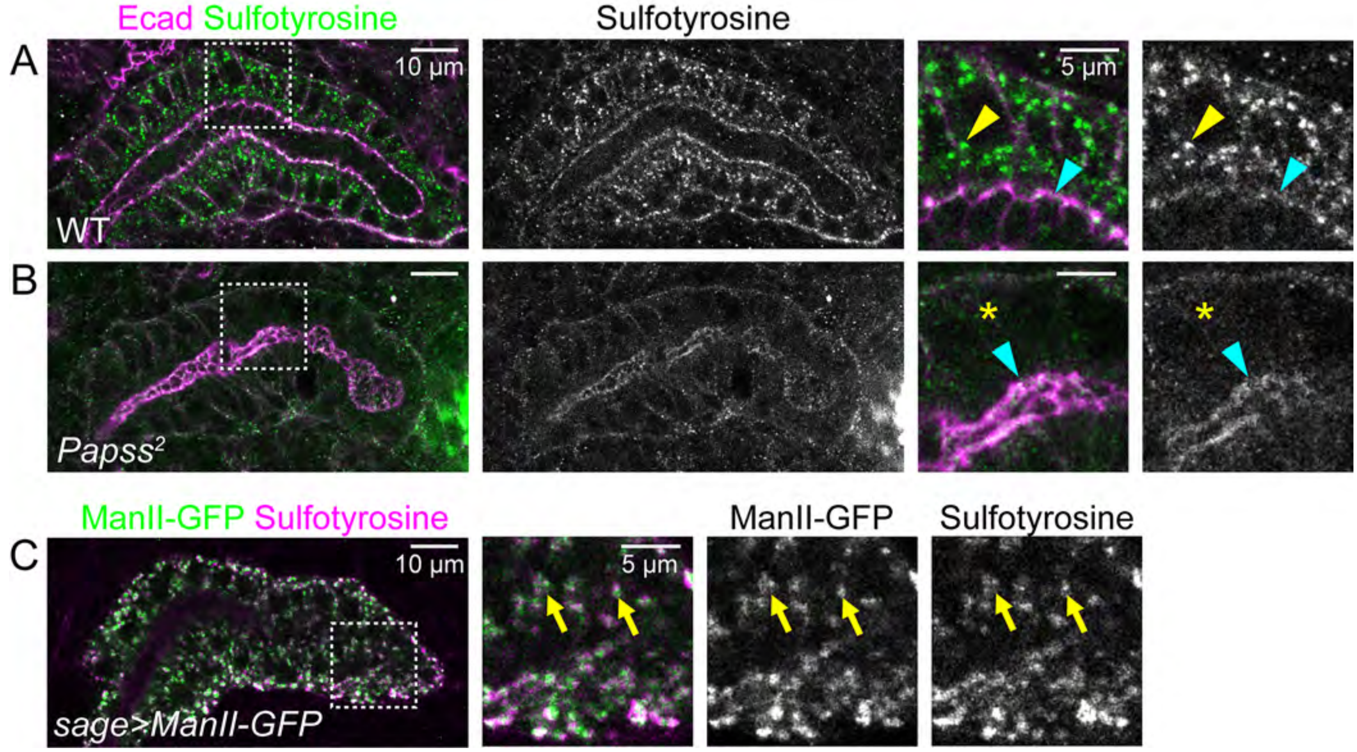


**Figure 1-figure supplement 2. Sulfotyrosine signals are reduced in *Papss* mutants.** (A, B) Confocal images of SGs immunostained for sulfotyrosine and Ecad in stage 16 WT (A) and *Papss* mutant embryos. Magnified images are shown for boxed regions. Yellow arrowheads, sulfotyrosine-positive cytoplasmic puncta in WT SG cells. Cyan arrowheads, sulfotyrosine at the apical membrane. Asterisks, almost no sulfotyrosine puncta are present in the cytoplasm in *Papss* mutants. (C) Confocal images of stage 16 SGs immunostained for GFP and sulfotyrosine. Arrows, ManII-GFP and Sulfotyrosine puncta co-localize.

Figure 1-figure supplement 3

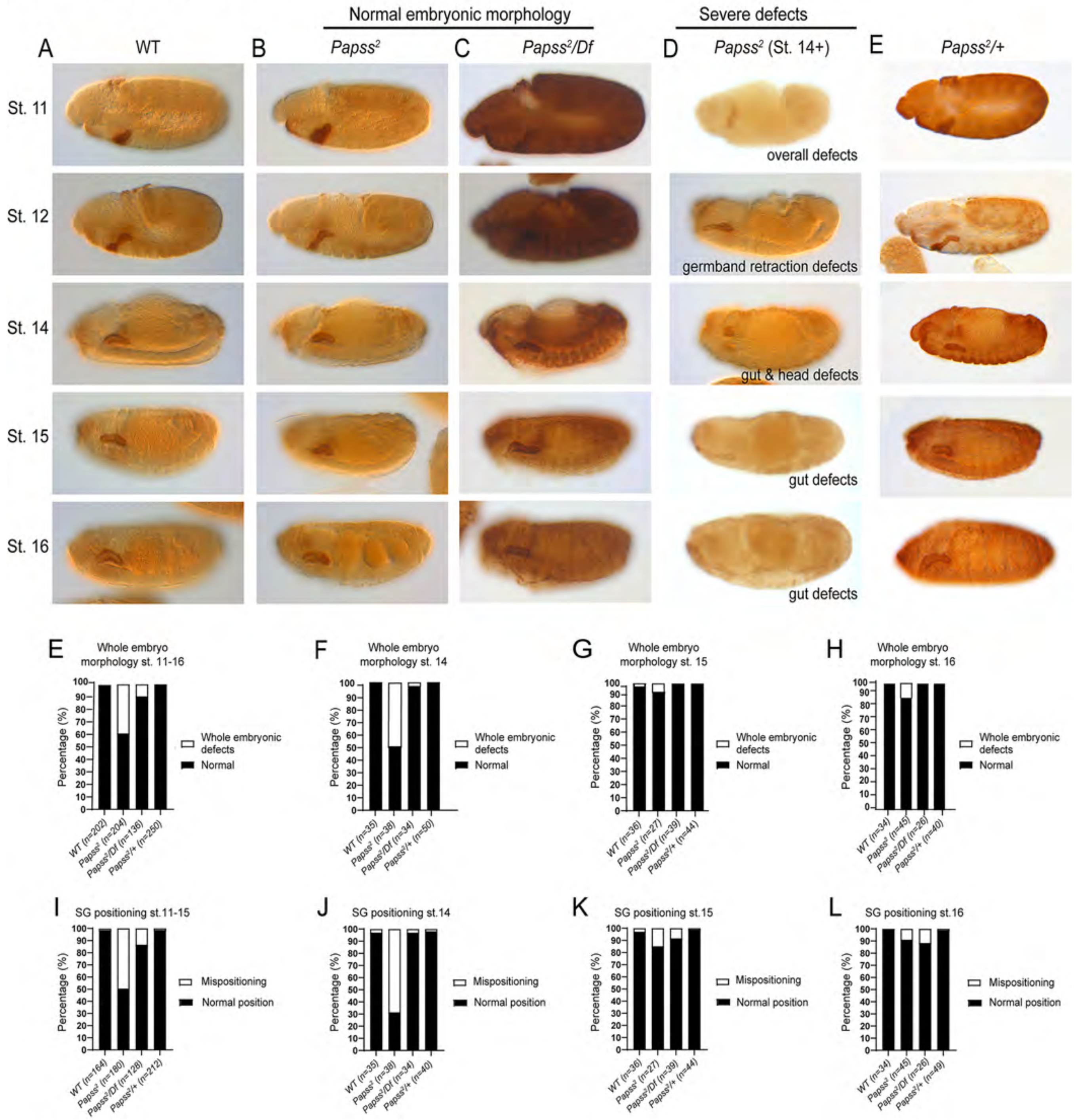


**Figure 1-figure supplement 3. Some *Papss* mutant embryos show defective SG formation and whole embryonic defects.** (A-D) Confocal images of stages 11-16 embryos immunostained using CrebA. (A) WT. (B) *Papss*, normal embryonic morphology. (C) *Papss*/*Df*, normal embryonic morphology. (D) *Papss*, severely defective whole embryo morphology. (E, F) Quantification of the percentage of embryos with whole embryonic defects. (E) Quantification of combined stages 11-16. (F) Quantification of individual stages for stages 14-16. (G-H) Quantification of the percentage of embryos with SG invagination defects/mispositioned SG. (G) Quantification of combined stages 12-16. (H) Quantification of individual stages for stages 14-16.

Figure 1-figure supplement 4

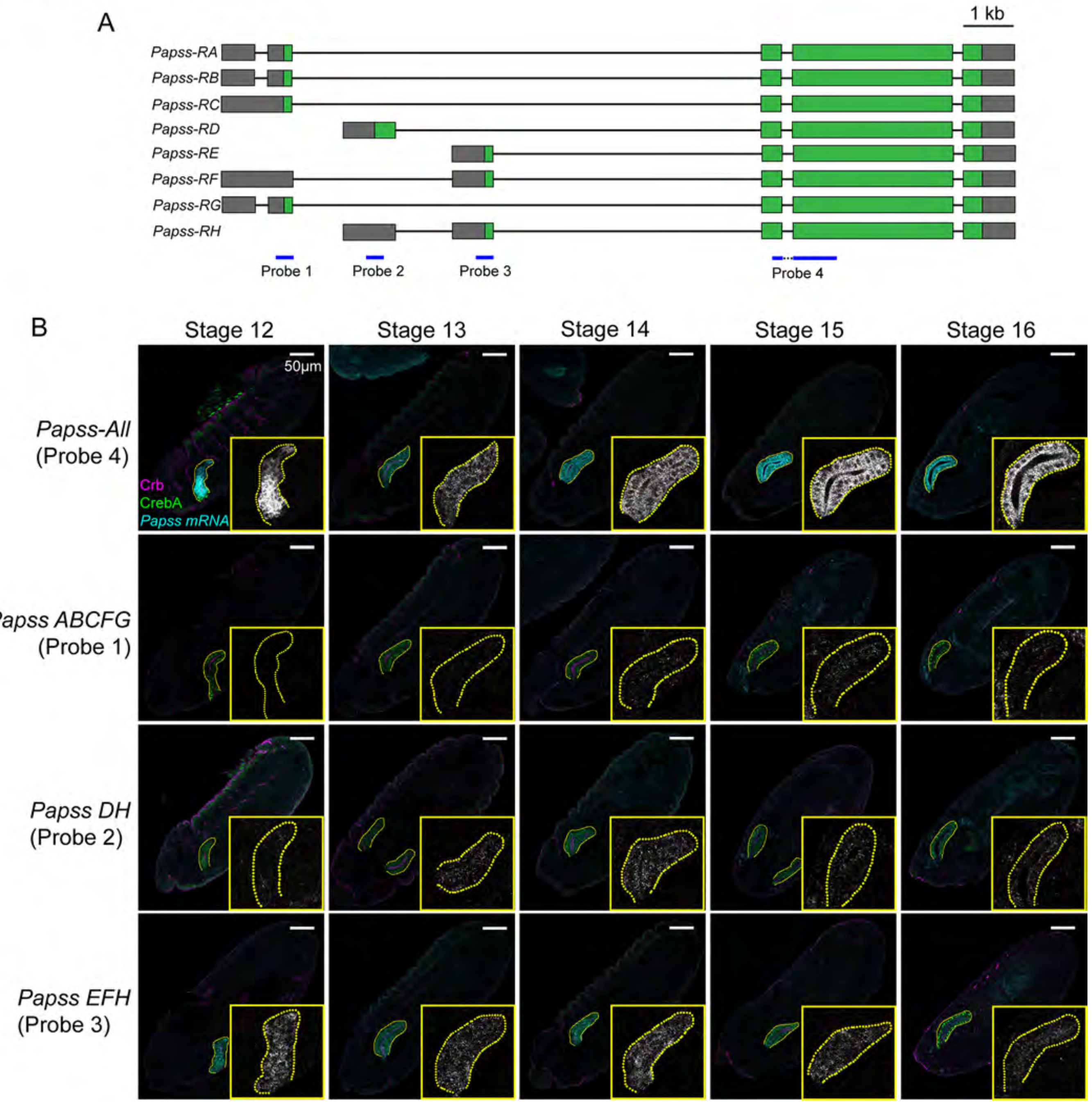


**Figure 1-figure supplement 4. Several splice forms of *Papss* are expressed in the SG.** (A) Eight annotated splice forms of *Papss* are reported by FlyBase. The blue bars indicate the regions used to make anti-sense probes that recognize different subsets of splice forms. Green and gray boxes indicate coding and non-coding exons, respectively. Thin lines indicate introns. (B) Fluorescence in situ hybridization with probes 1-4 (cyan). SGs are marked with CrebA (green) and Crb (magenta). Higher magnification of the mRNA signals within the SG are shown in the insets. Yellow dotted lines indicate the SG boundary.

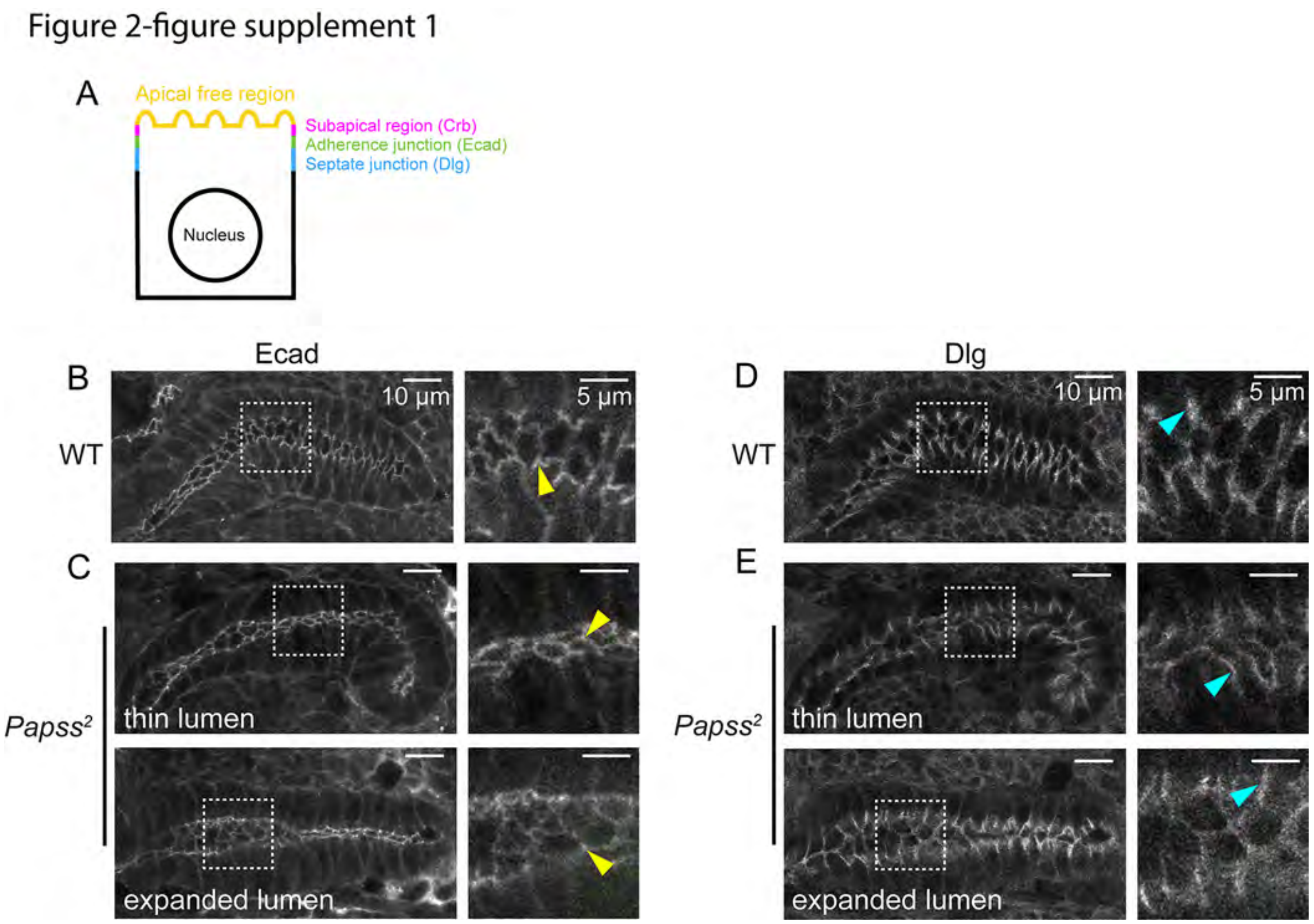


**Figure 2-figure supplement 1. Adherens junctions (AJs) and septate junctions (SJs) are intact in *Papss* mutants.** (A) Cartoon showing the subapical and junctional regions in the SG epithelial cell from the lateral view and localization of the key protein markers Crb, Ecad, and Dlg. (B, C) Confocal images of stage 16 SGs immunostained for Ecad. Magnified images are shown for boxed regions. Yellow asterisks, localization of Ecad to AJs in both WT (B) and *Papss* mutant (C) SGs. (D, E) Confocal images of stage 16 SGs immunostained for Dlg. Images show the luminal surface view. Cyan asterisks, localization of Dlg at SJs in both WT (D) and *Papss* mutant (E) SGs. SJs in *Papss* mutants occasionally appear slightly longer in the expanded luminal region compared to WT or the thin luminal region of *Papss* mutants, possibly due to stretching of cells with lumen expansion.

Figure 2-figure supplement 2

A WT_No.1

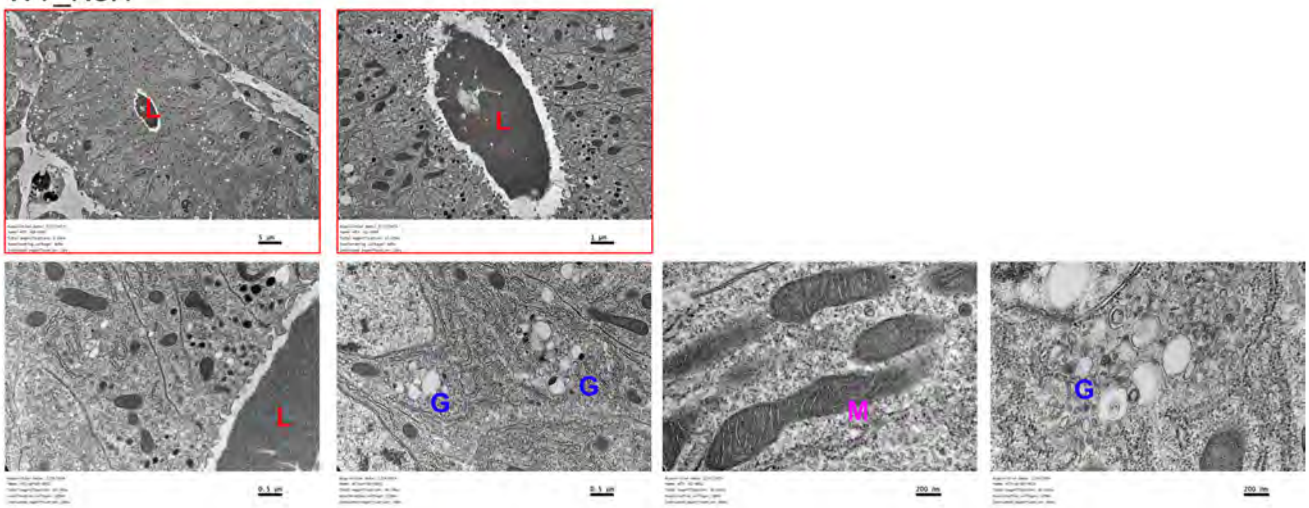


WT_No.2

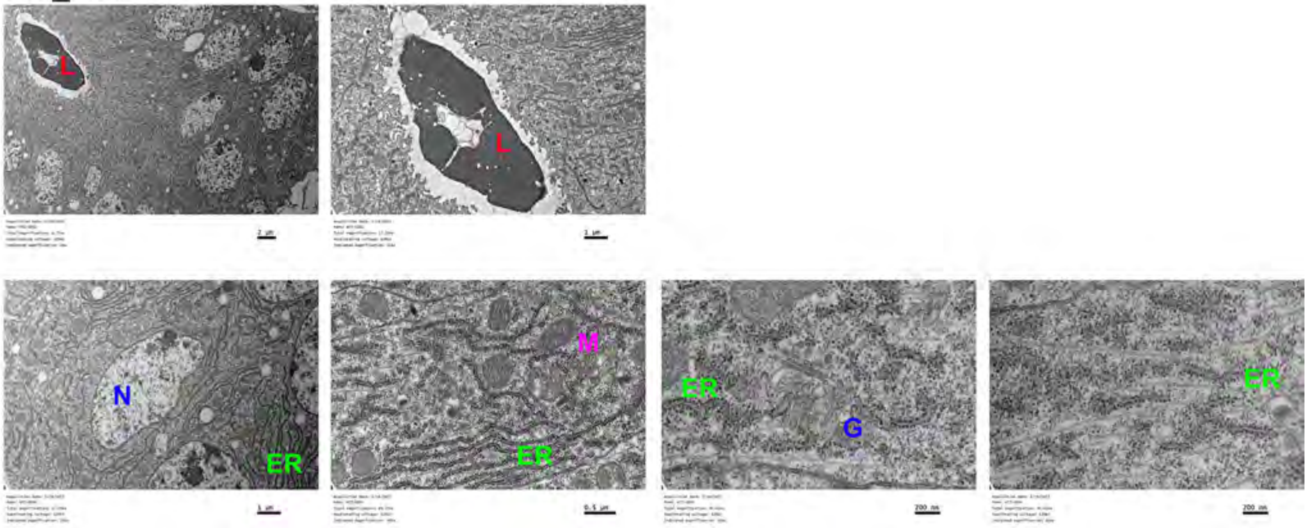

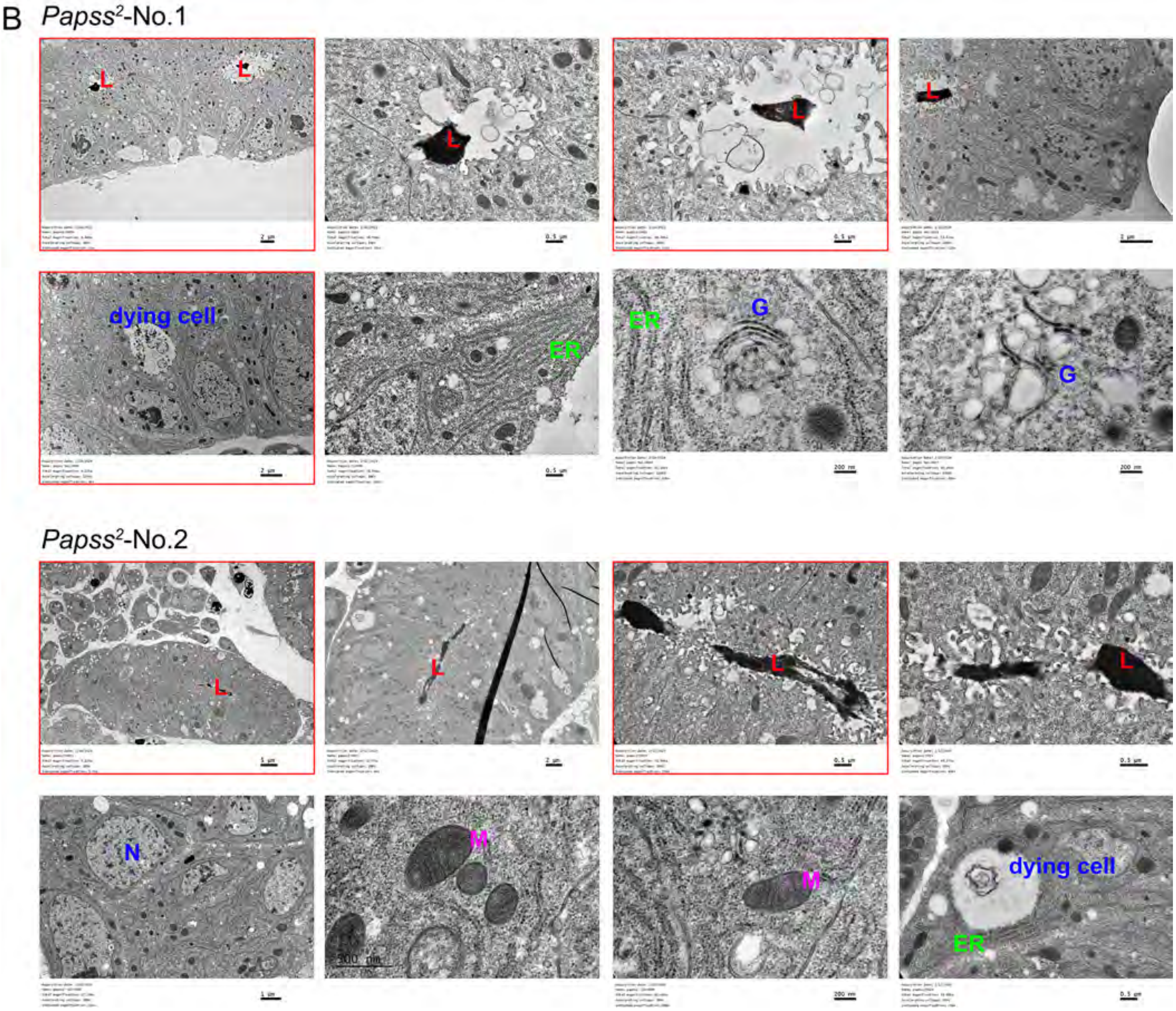


**Figure 2-figure supplement 2. Original, raw TEM images of WT and *Papss* mutant SGs.** Raw TEM images of two WT (A) and two *Papss* mutant (B) SGs at various magnifications. The images shown in Figures 2-4 are outlined in red. L, lumen. G, Golgi, M, mitochondria, ER, endoplasmic reticulum, N, nucleus.

Figure 3-figure supplement 1

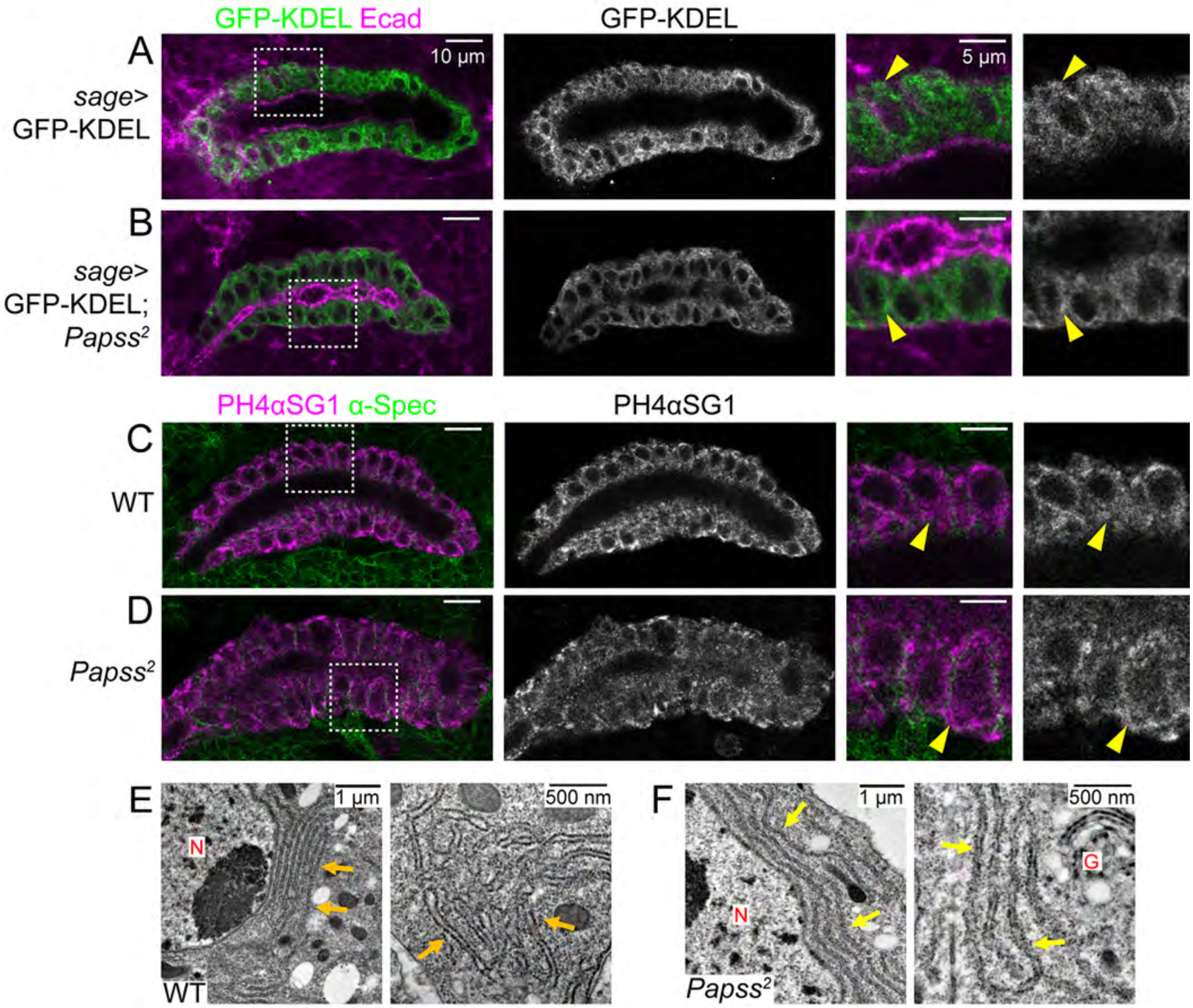


**Figure 3-figure supplement 1. The endoplasmic reticulum (ER) is intact in *Papss* mutants.** (A, B) Confocal images of stage 16 SGs immunostained for GFP (for GFP-KDEL) and Ecad. Magnified images are shown for the boxed regions. Yellow arrowheads, KDEL signals in the control (A) and *Papss* mutant (B) SGs. (C, D) Confocal images of stage 16 SGs immunostained for PH4αSG1 and α-Spectrin. Magnified images are shown for boxed regions. Yellow arrows, PH4αSG1 signals in the ER in WT (C) and *Papss* mutant (D) SGs. (E, F) TEM images of SG epithelial cells. N indicates the nucleus. G indicates the Golgi. Yellow arrows indicates the ER.

#### Figure 3-figure supplement 2

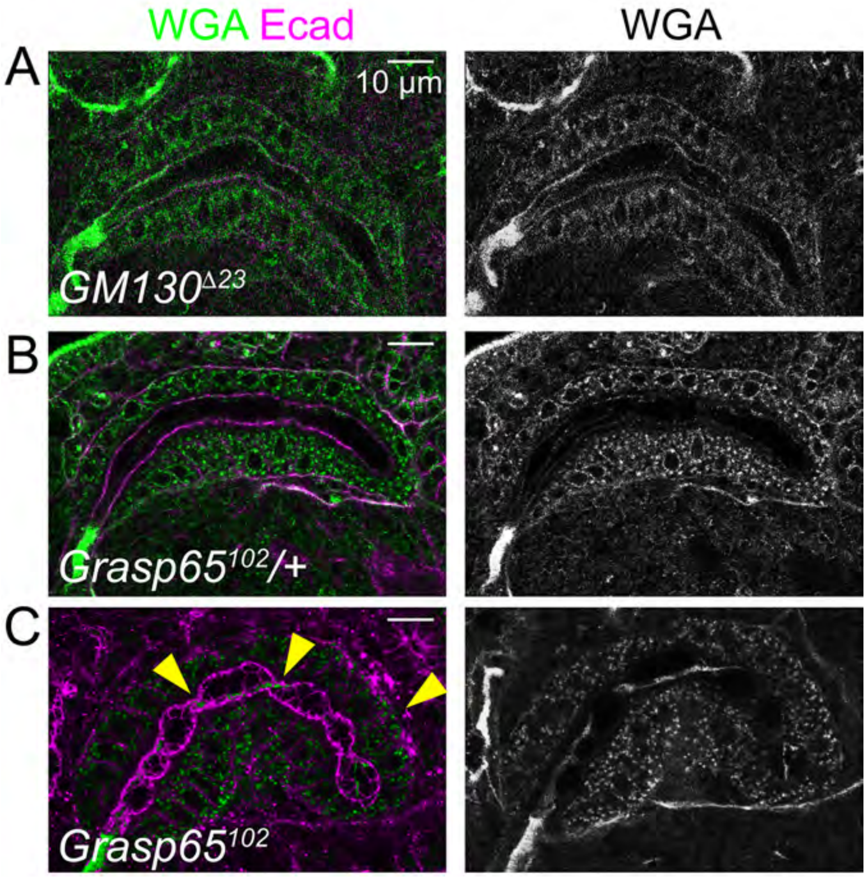


**Figure 3-figure supplement 2. Mutants of the Golgi component Grasp65 show an irregular SG lumen phenotype.** (A-C) Confocal images of stage 16 SGs immunostained for Ecad and WGA. Yellow arrowheads in C, constrictions in the SG lumen of the *Grasp65* mutant embryo.

#### Figure 3-figure supplement 3

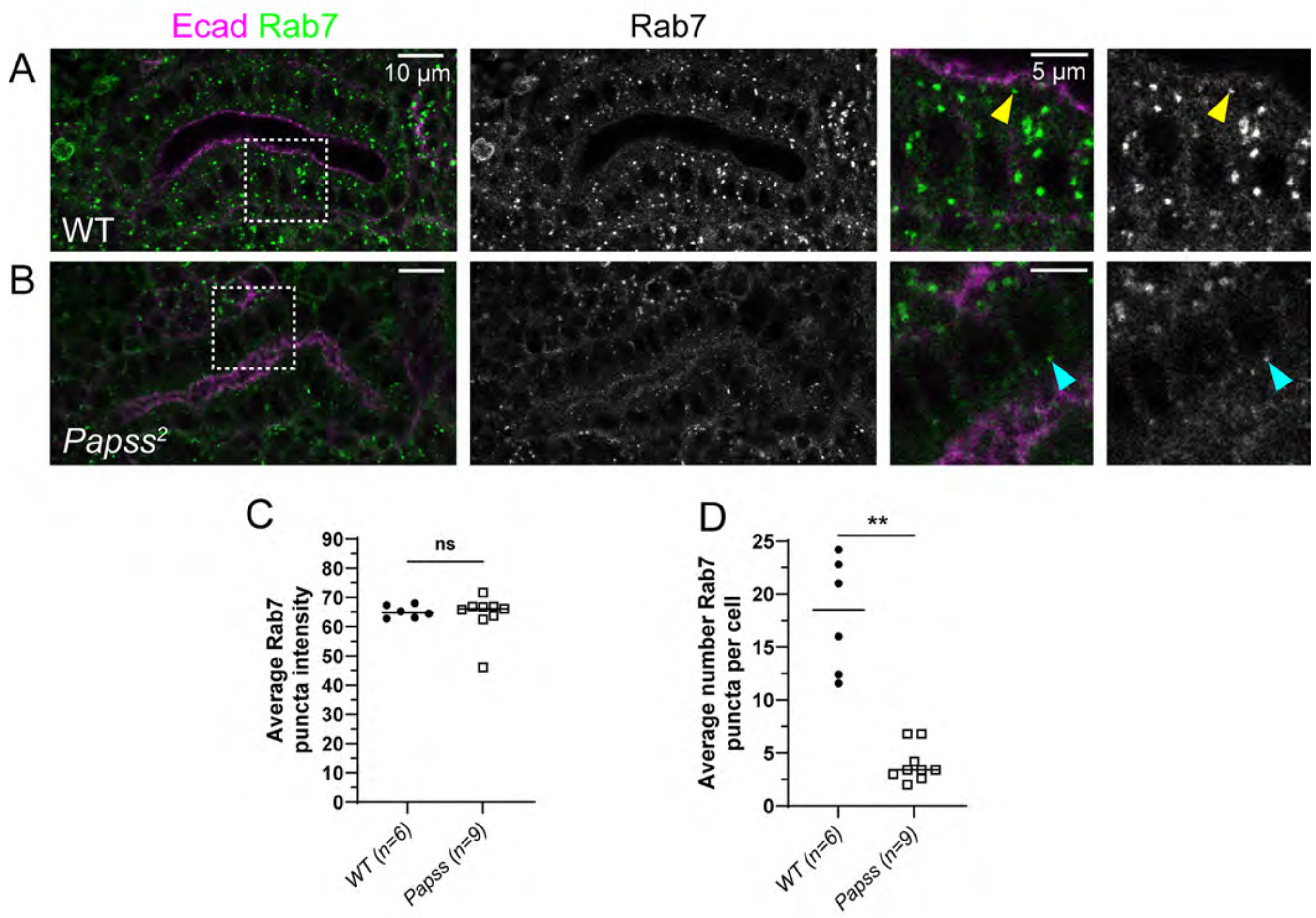


**Figure 3-figure supplement 3. Rab7 punctate number is decreased in *Papss* mutants.** (A) Confocal images of stage 16 SGs immunostained for Rab7 and Ecad. Magnified images are shown for boxed regions. Yellow arrowheads in A, Rab7 signals in WT. Cyan arrowheads in B, Rab7 puncta are reduced in *Papss* mutants. (B) Quantification for the average mean gray value intensity of Rab7 puncta per sample. (C) Quantification of the average number of Rab7 puncta found in each cell for one sample. (B-C) Welch's t-test (**, p<0.01). Horizontal lines indicate mean values.

Figure 5-figure supplement 1

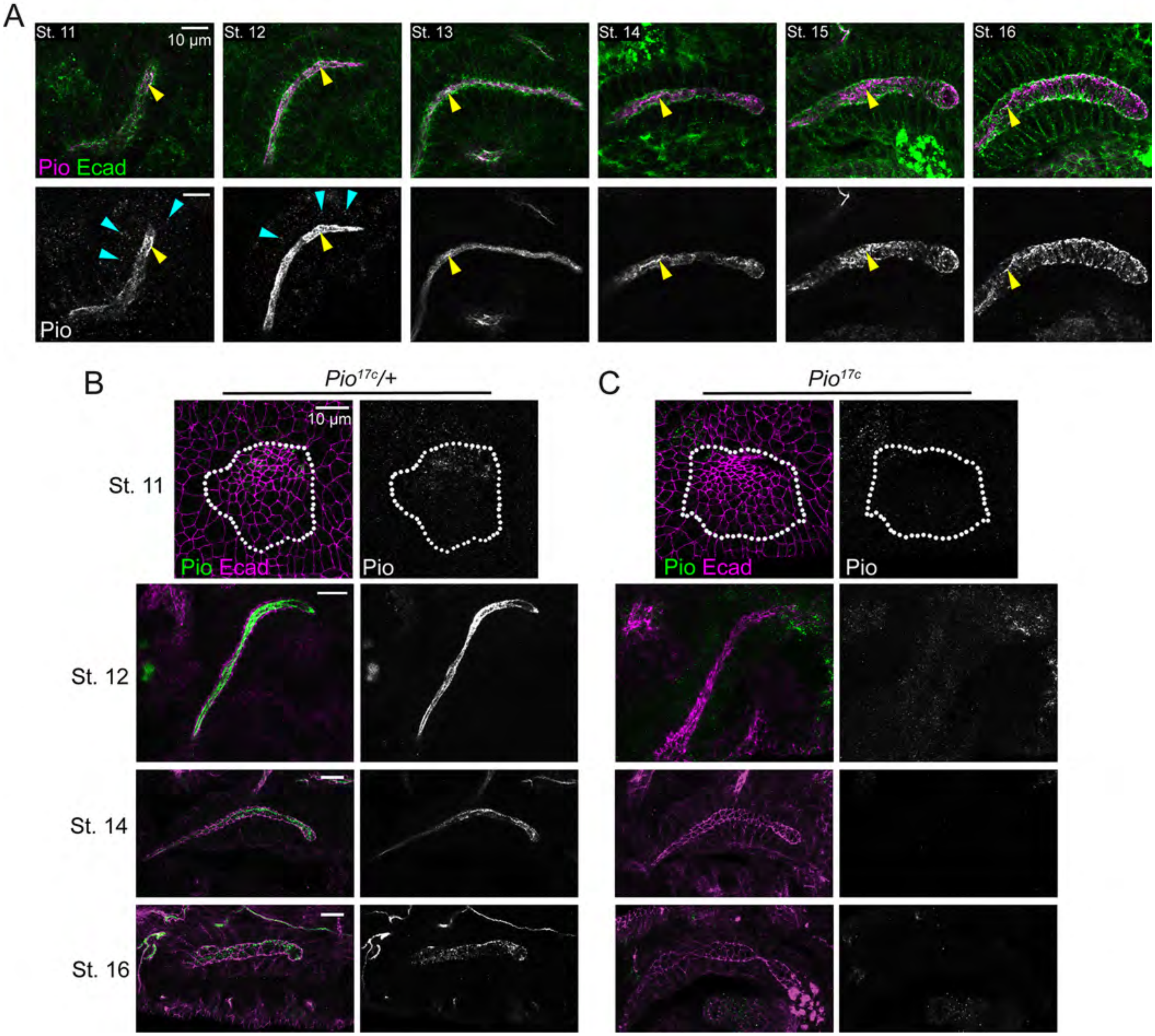


**Figure 5-figure supplement 1. Localization of Pio throughout SG development**. (A) Confocal images of stage 11-16 SGs immunostained for Pio and Ecad. Yellow arrowheads, Pio localizes to the SG lumen. Cyan arrowheads, Pio signals as cytoplasmic puncta. (B, C) Confocal images of stage 14 and 16 SGs immunostained for Pio and Ecad. Whereas filamentous luminal Pio signals are shown in control (B), no Pio signals are detected in *pio* null mutants (C).

Figure 5-figure supplement 2

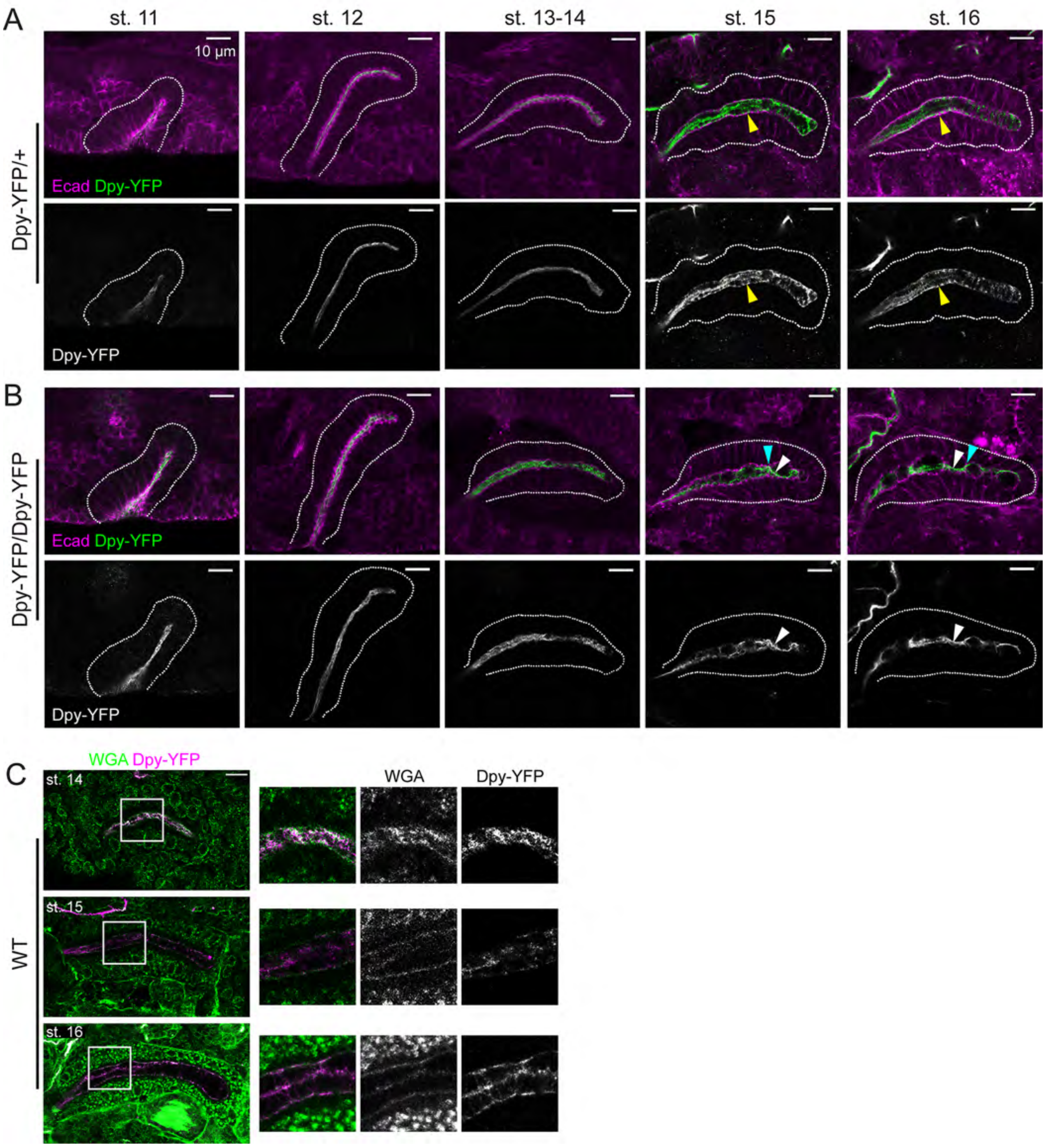


**Figure 5-figure supplement 2. Dpy localization through SG morphogenesis.** (A-B) Confocal images of stage 11-16 SGs immunostained using GFP and Ecad. (A) Dpy-YFP heterozygous samples. Yellow arrowhead, Dpy is localized in the SG lumen in an organized, filametous meshwork in Dpy-YFP/+. (B)

Dpy-YFP homozygous samples. Cyan arrowheads, the SG lumen shows sites of constriction in Dpy-YFP/Dpy-YFP. White arrowheads, Dpy organization is irregular in the SG lumen of Dpy-YFP homozygous embryos. (C) Stage 16 Dpy-YFP/+ SG stained for WGA. Luminal filamentous Dpy-YFP signals colocalize with WGA signals.

Figure 5-figure supplement 3

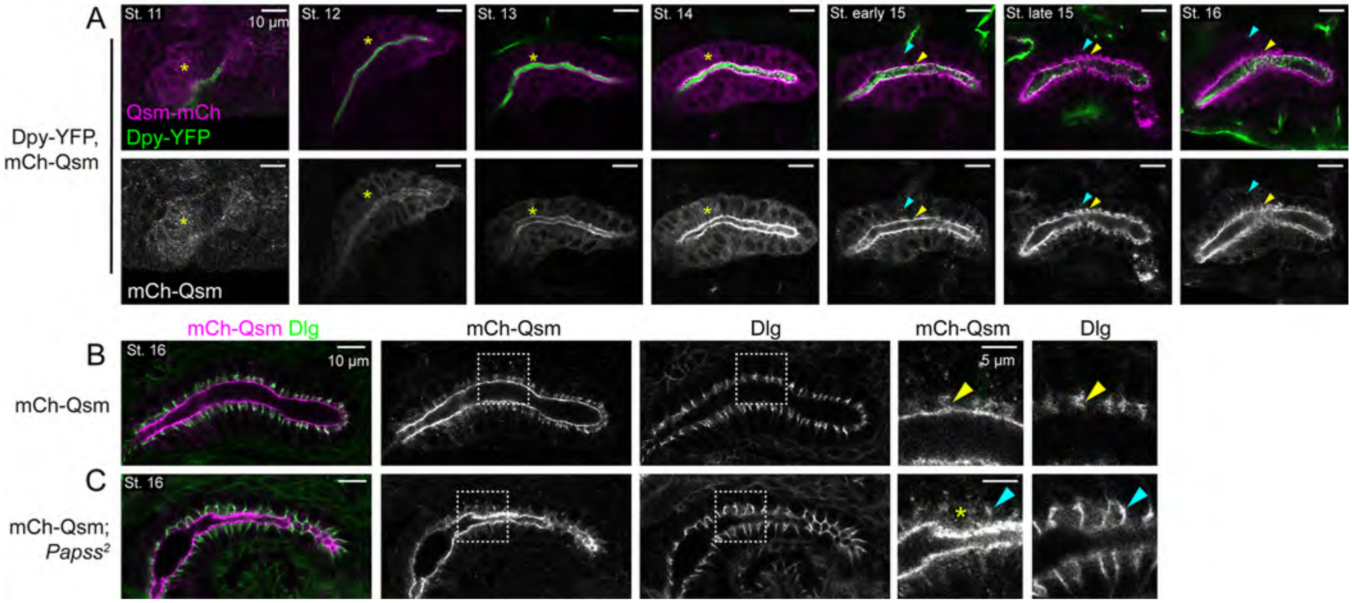


**Figure 5-figure supplement 3. Qsm localization patterns are not significantly affected in *Papss* mutants.** (A) Confocal images of stage 11-16 SGs immunostained for GFP (for Dpy-YFP) and mCh (for mCh-Qsm). At stages 11-14, mCh-Qsm localized throughout the cytoplasm of SG cells, with increased localization at the apical membrane. At late stage 14/early stage 15, the uniform cytoplasmic signals of mCh-Qsm began to decrease, and mCh-Qsm-positive vesicles formed near the apical domain and coalesced around SJs and the apical membrane. A few mCh-Qsm-positive vesicles were also observed in the basal region of SG cells. At stage 16, mCh-Qsm signals decreased in vesicles and instead increased at the apical membrane and were clearly detected at SJs, colocalizing with Dlg. Yellow asterisks, mCh-Qsm signals in the cytoplasm of SG cells. Yellow arrowheads, mCh-Qsm shows increased localization at the apical membrane and septate junctions in the late-stage SG. Cyan arrowheads, mCh-Qsm-positive puncta in the cytoplasm increase at early stage 15 in the SG. (B, C) Confocal images of stage 16 SGs immunostained for mCh and Dlg. Enlarged images are shown for boxed regions. mCh-Qsm partially colocalizes with Dlg in control (yellow arrowheads) and *Papss* mutant (cyan arrowheads) SGs. Yellow asterisks, dispersed mCh-Qsm signals near the apical membrane.

Figure 6-figure supplement 1

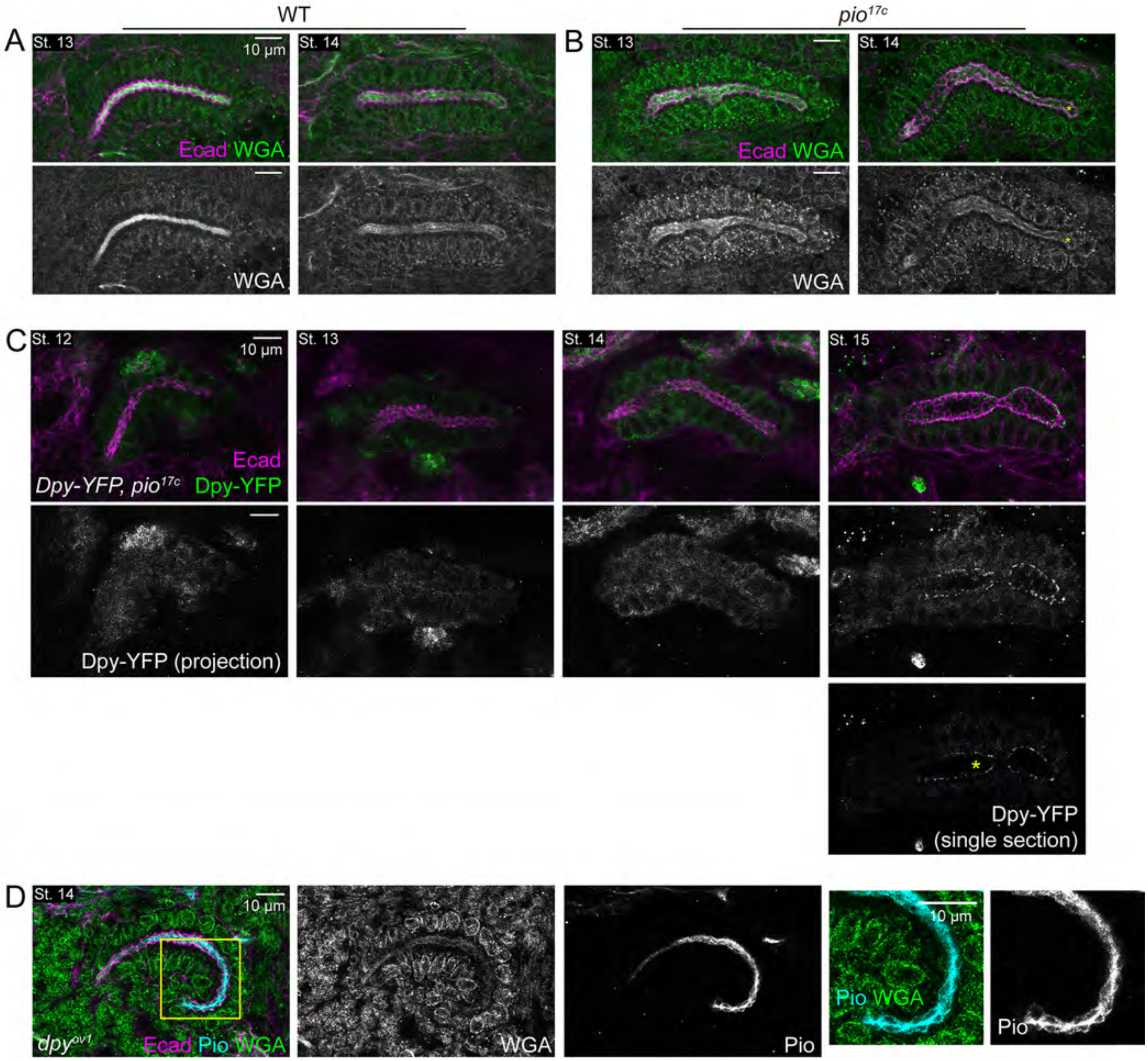


**Figure 6-figure supplement 1. Loss of *pio* affects Dpy localization patterns, and vice versa, in the SG**. (A, B) Confocal images of SGs immunolabeled for Ecad and WGA at stages 13 and 14 in WT (A) and *pio* mutant (B) embryos. (C) Confocal images of *pio* mutant SGs from stages 12-15 immunostained for Ecad and GFP (for Dpy-YFP). Projection images contain merged z-sections, from the apical surface of the SG cells to midway through the SG. A single section from stage 16 at the approximate midpoint of the lumen width. (D) Confocal images of *dpy* mutant SG at stage 14 immunostained for Ecad, Pio and WGA. Yellow boxes, area of higher magnification.
